# Efficiency of biological and chemical inducers for controlling Septoria tritici leaf blotch (STB) on wheat (*Triticum aestivum* L.)

**DOI:** 10.1101/2020.02.06.936583

**Authors:** Fares Bellameche, Chiara Pedrazzini, Brigitte Mauch-Mani, Fabio Mascher

## Abstract

The hemibiotrophic fungus *Zymoseptoria tritici* is the causative agent of Septoria tritici leaf blotch (STB) disease of wheat (*Triticum aestivum* L.), the economically most damaging disease of wheat in Europe. Today, ecofriendly plant protection methods compatible with sustainable agriculture are strongly desirable. Here, we applied the two chemical inducers β-aminobutyric acid (BABA) and benzo-(1,2,3)-thiadiazole-7-carbothioic acid S-methyl ester (BTH) and the two biotic inducers *Pseudomonas protegens* CHA0 (CHA0) and *P. chlororaphis* PCL1391 (PCL) on the roots of wheat seedlings in order to test their ability to induce resistance against STB. At 21 days after inoculation, only plants treated with BABA showed a smaller area covered by lesions and less pycnidia compared to the untreated control plants. We evaluated spore germination and fungal development on inoculated wheat leaves at early infection stages using calcofluor white staining. Overall, spores of *Z. tritici* germinated less on plants soil-drenched with BABA and BTH and their hyphal growth was significantly delayed. On the contrary, CHA0 and PCL seed treatments did not affect fungal growth in wheat leaves. In conclusion, BABA efficiently enhanced plant resistance to *Z. tritici*, BTH delayed fungal development at early stages while the two biotic inducers did not enhance resistance against STB disease.

## Introduction

The fungus *Zymoseptoria tritici* causes Septoria tritici blotch (STB) on wheat. *Z. tritici* is considered the most damaging wheat pathogen in Europe, mainly because of the suitable climatic conditions (Jørgensen et al. 2014). During severe epidemics, up to 50% of losses have been registered in a field planted with wheat cultivars susceptible to STB (Fones and Gurr 2015). Currently, no genetic resistance source in wheat cultivars is known to confer full resistance against STB. For control, farmers rely on the use of cultivars with partial resistance and on conventional fungicides (Schaad et al. 2019; Torriani et al. 2015). Yet, the durability of chemical control may remain ineffective in the field because *Z. tritici* displays a high capacity of adaptation and often succeeds in developing new resistances to fungicides and the overcoming of the host resistance (Cowger et al. 2000; Cheval et al. 2017). As chemical fungicides can have a negative impact on human and animal health and the environment (Aktar et al. 2009; Berny 2007), we decided to evaluate a more sustainable approach to combat this disease in the form of induced resistance.

One possibility of inducing disease resistance is the use of plant-growth promoting rhizobacteria (PGPR) and numerous cases of a successfully use of PGPR to improve plant nutrition and/or help plants overcome biotic or abiotic stresses have been documented (Chaudhary and Shukla 2019; Vurukonda et al. 2016; Dimkpa et al. 2009). These studies mainly involved the genus *Pseudomonas* commonly found among the predominant genera present in the rhizosphere and also on and in the root system of wheat plants (Yoshida et al. 2012). Many of these Pseudomonads are well-characterized PGPR and are able to exert plant-beneficial functions, including the suppression of plant diseases and stimulation of plant defences (Vacheron et al. 2013; Mauchline and Malone 2017). A subgroup including the species *P. protegens* and *P. chlororaphis* has been widely studied (Haas and Défago 2005; Mercado-Blanco and Bakker 2007; Hol et al. 2013; Vacheron et al. 2013). The strain *P. protegens* CHA0 (CHA0) is naturally suppressive to black root rot in tobacco (Stutz et al. 1986). It has been reported as a potential bacterial antagonist to control plant diseases (Hase et al. 2000; Ramette et al. 2011), and also for its capacity to induce resistance in dicotyledonous plants (Maurhofer et al. 1994; Iavicoli et al. 2003), as well as in monocots (Henkes et al. 2011; Sari et al. 2008). The strain *P. chlororaphis* PCL1391 (PCL) was first isolated from the rhizosphere of tomato, showing a high antagonistic activity against *Fusarium oxysporum*, the causal agent of tomato root rot (Chin-A-Woeng et al. 1998). The capacity of *P. chlororaphis* PCL1391 to protect plants against different other attackers has also been documented (Imperiali et al. 2017; Bardas et al. 2009; Flury et al. 2017).

Specific chemicals too have the capacity to enhance plant resistance to disease. For instance, synthetic compounds such as benzo[1,2,3]thiadiazolea-7-carbothionic acid-*S*-methyl ester (BTH, also called acibenzolar-S-methyl) and β-aminobutyric acid (BABA) have been reported to induce resistance in plants against a wide range of microbial pathogens without possessing direct antimicrobial activity (Görlach et al. 1996; Jakab et al. 2001; Justyna and Ewa 2013; Karthikeyan and Gnanamanickam 2011). The plant activator BTH is a functional analogue of salicylic acid and was one of the first chemical compounds shown to enhance activation of several defense responses against major fungal and bacterial pathogens in various crops including wheat (Iriti et al. 2004; Karthikeyan and Gnanamanickam 2011; Soleimani and Kirk 2012; Görlach et al. 1996; Vallad and Goodman 2004). BABA induces resistance in a wide range of economically important crop species and against a broad spectrum of pathogens including nematodes, virus, bacteria, oomycetes and fungi (Jakab et al. 2001; Barilli et al. 2010; Porat et al. 2003; Amzalek and Cohen 2007). Expression of BABA-induced resistance coincides with a faster and stronger defence response following pathogen attack, a phenomenon that has been termed priming (Cohen et al. 2016; Balmer et al. 2015). Even though BABA, BTH, CHA0 and PCL have been tested in many different pathosystems, to our knowledge, they have never been tested in wheat against STB.

*Z. tritici* is a filamentous fungal pathogen having the particularity of being hemibiotrophic, with two distinct phases of infection. Following inoculation onto the leaf surface by rain splash, spores germinate (Kema et al. 1996), giving rise to a single hypha. The hypha invades leaf tissues mainly through stomata (Rohel et al. 2001; Palmer and Skinner 2002) growing slowly in the apoplastic space between mesophyll cells, typically during up to 9–11 days (Kema et al. 1996; Shetty et al. 2003). Here the fungus survives by assimilating nutrients in solution in the apoplast (Shetty et al. 2003; Keon et al. 2007; O’Driscoll et al. 2014). This has also been referred to as ‘biotrophic’. During this first phase, no symptoms are visible. At a later time point, notably 12 to 20 days after penetration, the plant shows the first symptoms on the leaves. The appearance of symptoms coincides with the passage of the fungus to the ‘necrotrophic’ phase of the fungus (Palmer and Skinner 2002; Shetty et al. 2003). The infection process ends with the appearance of pycnidia, and leaf symptoms generally appear as light green to yellow chlorotic spots. As they enlarge, the lesions become brown and develop darker colored fruiting bodies (Ponomarenko et al. 2011).

In this study we aimed to assess the efficacy of the two chemical inducers (BABA and BTH) and of the two biological inducers (*P. protegens* CHA0 and *P. chlororaphis* PCL1391) as a preventive treatment on wheat seedlings against *Z. tritici*. The effect of the plant resistance inducers on fungal development was investigated at an early stage of infection. To exclude any direct inhibitory effect of the chemical inducers on fungal growth, we also assessed their direct antifungal activity towards *Z. tritici* by means of *in vitro* bioassays.

## Materials and methods

### Plant material and growth conditions

Experiments were carried out with the STB susceptible wheat variety Spluga (Agroscope/DSP). Prior to seeding, seeds were surface sterilized by rinsing with 70% ethanol and incubating for 5 minutes in 5 % bleach (sodium hypochlorite solution, Fisher Chemical, U.K.). Subsequently, the seeds were rinsed three times in sterile distilled water. The seeds were then pre-germinated for 3 to 4 days on humid filter paper (Filterkrepp Papier braun, E. Weber & Cie AG, 8157 Dielsdorf, Switzerland). We selected the seedlings with similar growth state and morphology to plant in 120 mL polypropylene tubes (Semadeni, 3072 Ostermundigen, Switzerland) filled with a standard potting mixture (peat/sand, 3:1, vol/vol). The plants grew in a growth chamber with the following conditions: 16 hours day at 22°C, 8 hours night at 18°C and an irradiance of 300 μmol m^−2^ s^−1^. The plants were watered as needed.

### Treatment with biological inducers

The biological inducers were used in the following trials were the rifampicin-resistant mutants CHA0-RIF of *Pseudomonas protegens* strain of CHA0 (Natsch et al. 1994), and PCL-RIF of strain *P. chlororaphis* PCL1391 (Chin-A-Woeng et al. 1998). The strains were routinely grown on solid King’s Medium B (KMB; *Pseudomonas* agar F, Merck KGaA, 64271 Darmstadt, Germany) supplemented with rifampicin 100 μg/mL at 24°C for 4 days. From this culture, a single colony was transferred to 100 mL of King’s liquid medium B (30g protease-peptone, 1.5g K_2_HPO_4_, 2.46 g MgSO_4_, 1.5g glycerol in 1 L distilled water) supplemented with 50 μg mL^−1^ rifampicin and incubated overnight incubation at 28°C with continuous shaking at 150 rpm. The resulting bacterial culture was centrifuged at 3700 rpm and washed twice with sterile 10mM MgSO_4_ solution. The final pellets were re-suspended in sterile distilled water and adjusted to 10^6^ cfu/mL (OD_600_ of 0.1) and used for seed inoculation. To this end, the wheat seeds were immerged into the bacterial suspension for 6 hours with shaking at 35-40 rpm at room temperature. Control seeds were soaked in distilled water for the same duration. Inoculated seeds underwent the pre-germination procedure as described above.

### Chemical inducer treatment

BTH formulated as BION^®^ 50 WG (50% active ingredient) was obtained from Syngenta (Basel, Switzerland) and BABA from Sigma-Aldrich (Buchs, Switzerland). BTH (2mM) and BABA (15mM), respectively, were dissolved in distilled water and 10 mL per growing tube were used as soil-drench 2 days before *Z. tritici* inoculation. Control plants were just watered with distilled water. The concentration of BTH used in this study was chosen according to Görlach et al. (1996). While, BABA at 15 mM was chosen as ideal concentration to induce resistance against leaf rust without any effect on plant growth (unpublished data).

### Fungal cultures and plant inoculation

*Z. tritici* isolate 3D7 (Zhan et al. 2002) was provided by Prof. Daniel Croll (University of Neuchâtel, Switzerland). The isolate was stored at −80°C in 50% glycerol. Stock cultures were cultivated on Yeast-Sucrose Agar (YSA; 10 g L^−1^ yeast extract, 10 g L^−1^ sucrose, 1.2% agar) supplemented with kanamycin (50 μg/mL). For inoculum preparation, the strain was cultured in liquid YSB amended with 50 μg/mL kanamycin and incubated for 8 days at 18°C under continuous shaking at 150 rpm. After incubation, the suspension was filtered through a sterile cheese cloth and rinsed with sterile distilled water. Prior to infection, the spore concentration was adjusted to 10^5^ spores/mL in distilled water using a haemocytometer. After adding 0.1% tween20 to the spore suspension, each plant was spray-inoculated until runoff. The plants were then maintained at 100% relative humidity for 48 hours. After this, the plants were placed in a growth chamber as described above.

### Infection quantification

At 21 days after inoculation (dai), symptoms on wheat plants were quantified as described by Stewart et al. (2016). Briefly, the third leaf of each plant was excised, fixed on a sheet of paper and immediately scanned at 1.200 dpi (Epson perfection, V370 PHOTO). The leaf surface covered with pycnidia, lesions or leaf necrosis was measured using an automated image analyses macro for the software ImageJ version 1.x (Schneider et al. 2012). The disease severity was the expressed as percentage of leaf area covered by lesions (PLACL).

### *In planta* fungal growth

Monitoring of spore germination and hyphal growth of *Z. tritici* on the leaf surface was performed using Calcofluor White (Sigma-Aldrich, Germany) staining according to Siah et al. (2010). Briefly, third leaf segments (4 cm) from three randomly selected replicates of each treatment were harvested at 24, 48 and 120 hours after inoculation (hai) and immersed for 5 minutes in 0.1% (*w/v*) Calcofluor White solution prepared in 0.1 M Tris-HCl buffer pH 8.5. After washing with sterile distilled water, the leaf segments were dried in darkness at room temperature. After covering with a cover slip, the preparations were examined under the epifluorescence microscope (Model E800; Nikon Instruments Europe, Badhoevedorp, The Netherlands) using excitation at 365 nm in combination with a 450 nm barrier filter and a dichroic mirror at 400 nm.

### *In vitro* antifungal assay

A potential direct effect of BTH and BABA on growth of *Z. tritici* was spectrophotometrically assessed in liquid YSB supplemented with kanamycin 50 μg/mL. Since BION contains additional ingredients that can influence the absorbance measurement, the active molecule Acibenzolar-S-methyl (Sigma-Aldrich, Germany) was used. BABA and BTH were first dissolved in distilled water and filter-sterilized with a 0.22-μm syringe filter (Millex GP, Millipore). Concentrations of 0, 0.02, 0.2 and 2 mM of BTH and 0, 0.15, 1.5 and 15 mM of BABA were tested. Aliquots of 40 mL culture medium were inoculated with 50 μL of fresh fungal spore suspension (10^5^ spores/mL) and placed at 18°C in the dark under continuous shaking at 150 rpm. Fungal growth was assessed daily by measuring the optical density at 405 nm.

### Statistical analyses

All experiments were repeated twice. Infection quantification was carried out in eight biological replicates. The germination of conidia *in planta* was observed in at least 50 spores on three independent replicates. For both assessments, PLACL and the germination class of conidia, comparisons between the treatments were carried out with one-way ANOVA. After ascertaining that the residues were normally distributed, significant differences between treatments were tested *post-hoc* using Tukey’s HSD test.

For the growth inhibition assay, the area under the growth curve was calculated for each BABA and BTH concentration in three independent replicates. Significant difference in response to dose-treatment were analysed by a Student’s *t*-test in comparison to the control (0 mM BABA or BTH). In all trials, significant differences were considered at *P* <0.05. All statistical analyses were conducted in R (R Core Team 2018).

## Results

### Plant response to *Z. tritici* after pre-treatment with resistance inducers

Symptoms on leaves were assessed on the third leaf, at 21 days after infection (Fig. 1a). Infected leaf tissue initially became chlorotic and later turned necrotic. In the untreated control, in the bacteria-treated plants and in the BTH treatment a large proportion of the leaves was necrotic and only a small part was alive (green). In opposite, plants treated with 15mM BABA presented less symptoms compared to the untreated control and leaves were green. The extension of the lesions (PLACL) was significantly lower in plants treated with BABA in comparison with the untreated control and the other pre-treatments (Fig. 1b). Similarly, the density of pycnidia was significantly reduced in BABA but not in the other treatments (Table 1).

**Table 1.** Pycnidia density per leaf (pycnidia/cm^2^) in H_2_O-treated control plants and CHA0-, BCL-, BABA- or BTH-treated plants. Pycnidia density was assessed on the third leaf of each biological replicate (n = 8).

| Treatment | Pycnidia/cm<sup>2</sup> |
| --- | --- |
| H<sub>2</sub>O | 97.79 ± 39.78<sup>a</sup> |
| CHA0 | 109.94 ± 25.04<sup>a</sup> |
| PCL | 112.39 ± 13.76<sup>a</sup> |
| BABA | 14.29 ± 19.81<sup>b</sup> |
| BTH | 88.48 ± 29<sup>ab</sup> |

**Fig 1.**
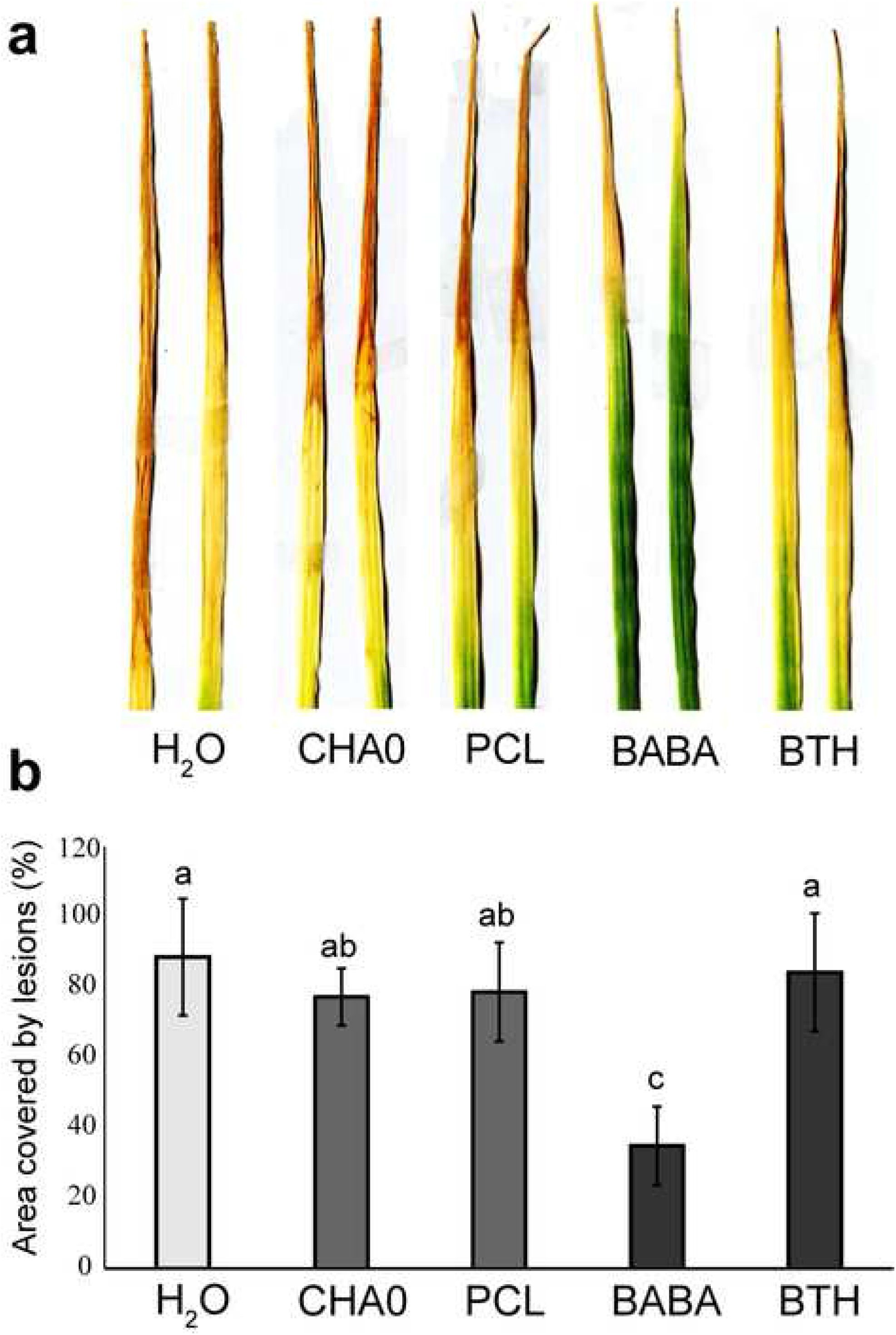
Response of wheat seedlings cv. Spluga to infection with *Z. tritici* after pre-treatment with H_2_O (control), CHA0, PCL, BABA and BTH, respectively. Symptoms were observed at 21 dai (A) and percentage of total leaf area covered by lesions (B) was obtained from scanned images analysed with an ImageJ macro. Error bars indicate the standard error for the average values of 8 replicates. Bars with the same letter are significantly not different at P < 0.05.

### Early effect of plant resistance inducers on *in planta* spore development

Spore germination and hyphal growth of *Z. tritici* on the leaf surface of wheat plants (cv. Spluga) and growth structures were counted at 24, 48 and 120 hai. To quantify the effect of resistance inducers during this observation period, four distinct developmental classes have been defined (Fig. 2a): class 1, spore non-germinated; class 2, geminated spore with a short germ tube; class 3, geminated spore with a well-developed germ tube; class 4, germinated spore with branched hyphae. Fig. 2b shows the percentage of these classes in each treatment. At 24 hai, fungal development displayed similar proportions of class 1 and class 2. Only in the control and in the CHA0 treatment a small proportion of class 3 structures (about 3%) were present. At 48 hai, the control presented about 45% of class 1, 42% of class 2 and about 13% of class 3 structures. The proportions in the bacterial treatments were similar to the control and statistically not different. Following chemical resistance induction, more spores were in class 1 and 2 compared to the control. While the proportions were statistically not different between BTH-treatment and the bacterial treatments, the BABA treatment presented a significantly higher number of class 1 structures compared to the bacterial CHA0 treatment.

**Fig 2.**
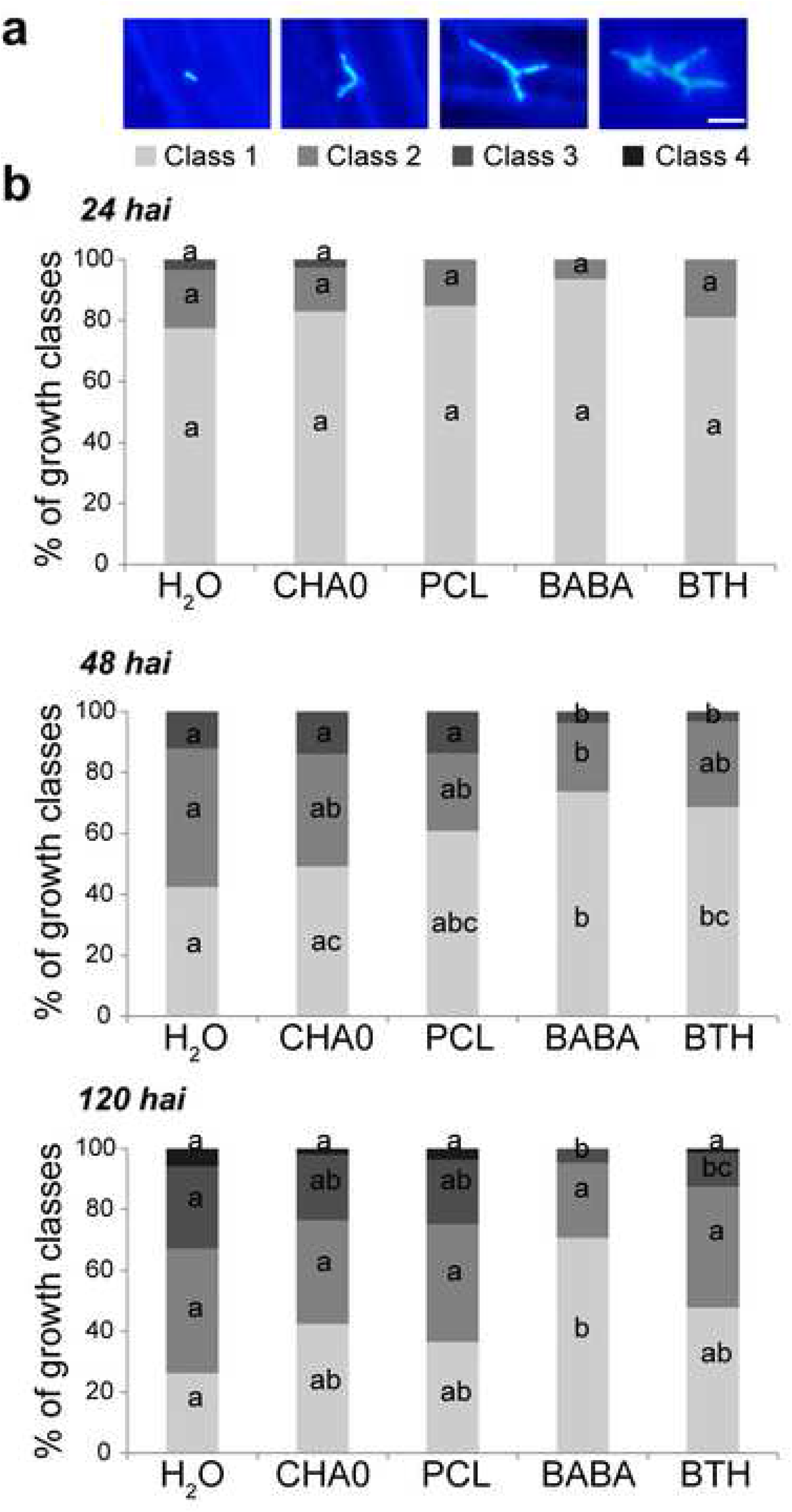
Effect of different plant resistance inducers on spore germination and hyphal growth of *Z. tritici* on leaves of wheat cv. Spluga. Four types of fungal developmental classes were defined (A): class 1, spore non-germinated; class 2, geminated spore with a short germ tube; class 3, geminated spore with a well-developed germ tube; class 4, germinated spore with branched hyphae. Scale bar = 10 μm. On 3 biological replicates, 50 spores were observed. The percentage of each class was assessed in each treatment at 24, 48 and 120 hai (B). Bars with the same letter are significantly not different at P < 0.05.

At 120 hai, a small proportion of class 4 (hyphae with branching) was present in the control and both bacterial treatments and BTH treatment but not in BABA-treated plants. This was also true for growth class 1. In BABA-treated plants, 70% of spores were in growth class 1. This was 25% higher than in BTH-and CHA0-treated plants. The number of spores in class 1 was not different between the CHA0, PCL, BTH and the control treatment. For class 2, the number of spores that produced a small germ tube did not differ between treatments. Yet, the number of spores with a well-developed germ tube (class 3) was not significantly different between the bacterial treatments and the control and the bacterial treatments and BTH-treated plants. This proportion was very small in BABA-treated and not different between BABA-and BTH-treated plants but highly significantly different between BABA / BTH treatments and the control.

### *In vitro* antifungal activity of BABA and BTH on *Z. tritici* growth

In order to test whether BABA or BTH have a direct inhibitory effect on fungal growth, *Z. tritici* was grown in YSB liquid medium amended with the two inducers. A dose-dependent inhibition was observed for both chemicals (Fig. 3). No antifungal activity was observed for all tested BTH concentrations. At the highest concentration (BTH 2 mM), *Z. tritici* growth was slightly inhibited (Fig. 3a) but no significant differences were observed. When BABA was added to the medium only the highest tested concentration (15 mM) led to a significant delay in fungal growth compared to the control without BABA (Fig. 3b).

**Fig 3.**
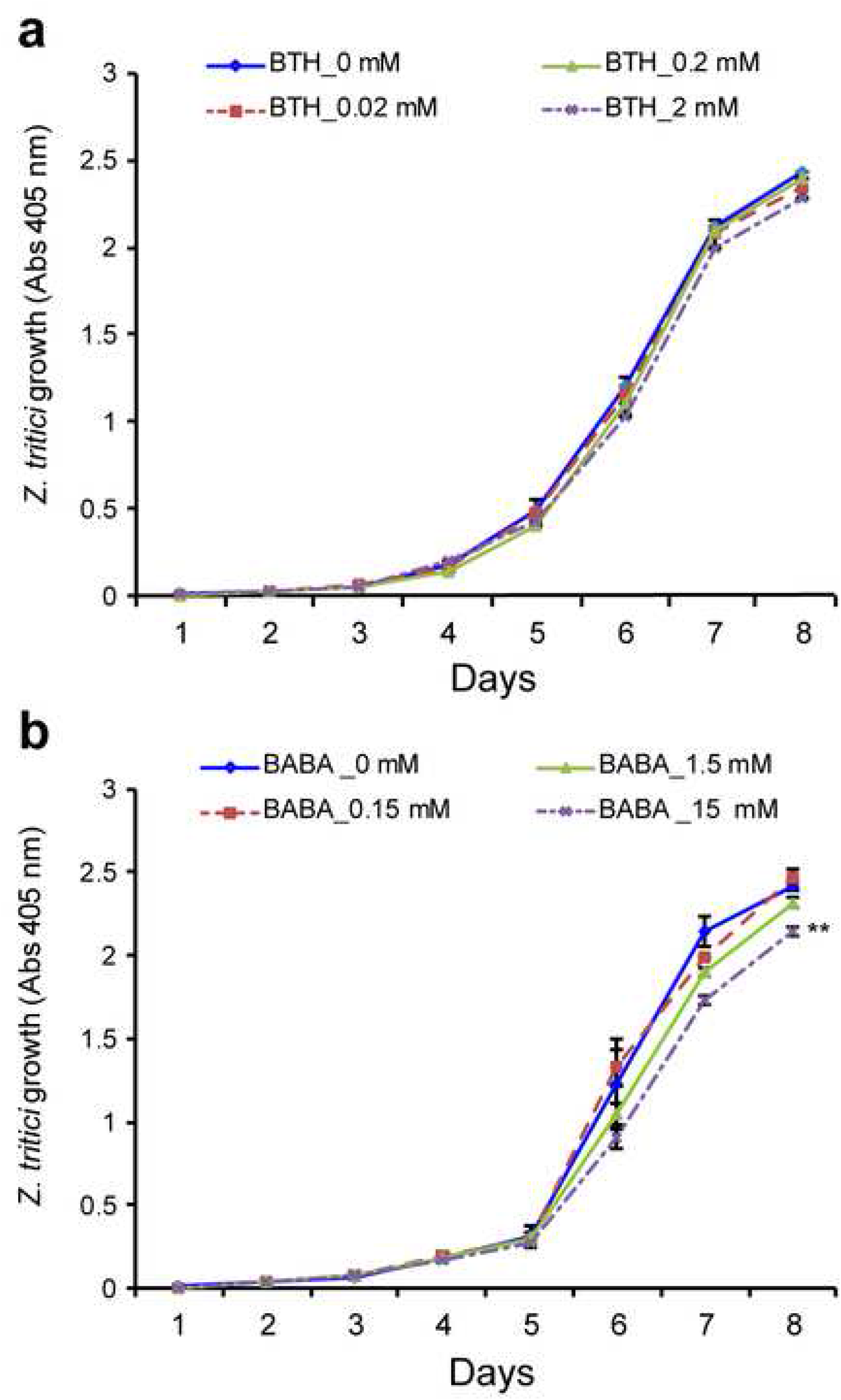
*In vitro* dose–response curves of *Z. tritici* to BABA (A) and BTH (B). Fungal growth was spectrophotometrically measured at 405 nm during 8 days in YSB amended with BABA 0.15, 1.5 and 15 mM or BTH 0.02, 0.2 and 2 mM. Error bars indicate the standard error for the average values of 3 replicates. Asterisks indicate significant differences in area under curves in response to dose-treatment determined by Student’s *t*-test: *P<.05; **P<.01; ***P<.001.

## Discussion

Plant resistance inducers are a promising alternative to control fungal disease (Wang and Zhou 2018; Chaudhary and Shukla 2019). Here, we report on the possibility to significantly reduce the severity of Septoria leaf blotch, caused by *Z. tritici* when BABA is applied as a preventive treatment in wheat.

General leaf symptoms of Septoria leaf blotch, such as chlorosis and necrosis, were assessed during 21 dai. Severe symptoms were observed in the untreated control as well as in BTH-, PCL- and CHA0-treated plants. Only BABA-treated plants showed a significantly lower percentage of leaf area covered by lesions (PLACL) and a significantly lower number of pycnidia.

To understand the response to infection with *Z. tritici* in wheat treated with resistance inducers, at the early stage, the fungal development was tracked by microscopy at 24, 48 and 120 hours post-infection. As expected, in BABA-treated plants, pathogen growth was significantly delayed. Therefore, BABA as a well-known priming agent, presumably activated a fast and robust response to fungal attack in the host plants (Zimmerli et al. 2000; Ton and Mauch-Mani 2004; van Hulten et al. 2006).

Exogenous application of BABA can inhibit development of disease directly or indirectly (Cohen et al. 2016). Since BABA is highly systemic, readily taken up by roots, and transported to the leaves (Cohen and Gisi 1994), it was not possible to conclude whether the observed resistance was direct or not. A potential direct fungicidal action by BABA on the growth of *Z. tritici* was excluded in the *in vitro* growth assay. Only a high concentration of BABA (15 mM) reduced fungal growth. Similar results showing that a high concentration of BABA exhibited a toxic effect on pathogen *in vitro* growth have been reported. Porat et al. (2003) observed that a very high concentration of BABA (100 mM) completely inhibited spore germination and mycelial growth of *Penicillium digitatum.* Similarly, the addition of BABA (50 to 200 mM) to the suspension culture of *Penicillium italicum* inhibited spore germination and germ tube elongation *in vitro* (Torriani et al. 2015). In another study, Fischer et al. (2009) showed that BABA inhibited mycelial growth and germination of *Botrytis cinerea* in a concentration-dependent manner, suggesting that direct antifungal effects of BABA may be associated with its concentration. In our study, low concentrations of BABA (0.15 and 1.5 mM) did not limit *Z. tritici* growth. It is important to note that the concentration of BABA inside wheat leaves at 2 and 6 days post application of 15 mM BABA to the roots were 16 and 6 µM respectively (Table S1), this is far below the *in vitro* inhibition concentration. Therefore, for our *in planta* assays, we postulate that BABA primes resistance mechanisms of the plant that inhibit the germination of *Z. tritici* in the wheat leaves.

As mentioned before, at 21 dai, BABA-treated plants showed less PLCAL and a lower number of pycnidia. This could be explained by results observed in the fungal growth assessment, where BABA treatment significantly limited *Z. tritici* growth. During the transition to the necrotrophic phase, the fungus releases cell wall-degrading enzymes such as β-1,4-endoxylanase, which have been shown to be correlated with symptom and sporulation levels of *Z. tritici* (Siah et al. 2010; Douaiher et al. 2007). Hence, the limitation in fungal growth may decrease the production of cell wall-degrading enzymes resulting in less PLCAL and a lower number of pycnidia.

In the initial infection phase (48 hai), BTH limited fungal development. However, the effect of BTH did not persist during the whole infection process. One hundred and twenty hours after inoculation, hyphal development on BTH-treated plants hardly differed from the non-treated controls. In addition, 21 dai, symptoms on plant leaves were similar to untreated plants. This suggests that BTH did not enhance resistance against *Z. tritici* in wheat seedlings. We suppose that the delay of spore germination observed during the initial infection phase, may be due to an indirect effect since none of the tested concentrations of BTH delayed or inhibited *Z. tritici* growth *in vitro*. Recently, Mejri et al. (2019) studied the protection efficacy of several salicylic acid conjugated derivatives on wheat against *Z. tritici*, and observed no correlation between direct fungicidal activity *in vitro* and protection of wheat plant.

Previous studies have shown that BTH enhances plant resistance to fungal pathogens by activating the systemic acquired resistance signal transduction pathway (Benhamou and Bélanger 1998; Liu et al. 2005; Azami-Sardooei et al. 2013; Abdel-Monaim et al. 2011). BTH treatment induces the accumulation of many transcripts that also accumulate during pathogen infection in Arabidopsis (Görlach et al. 1996; Lawton et al. 1996). In wheat, BTH can induce resistance to powdery mildew (*Blumeria graminis*), leaf rust (*Puccinia triticina*) and Septoria leaf spot and this resistance is accompanied by the induction of a number of wheat chemically induced (WCI) genes (Görlach et al. 1996). However, BTH did not provide resistance to Fusarium head blight in wheat (Yu and Muehlbauer 2001).

Neither CHA0 nor PCL induced plant resistance to *Z. tritici* infection in wheat Seed treatment by both rhizobacteria did not affect spore germination and hyphal growth in the early infection phase of *Z. tritici*. Moreover, symptoms on bacteria-treated plants were the same as in water-control plants. Following seed inoculation, both bacteria colonized the wheat roots to more than 10^5^ CFU/g of root fresh weight (data not shown). This degree of colonization provided effective plant protection in soils suppressive to take-all of wheat and barley caused by *Gaeumannomyces graminis* var. *tritici* (Weller et al. 2007), Fusarium wilt of pea mediated by *Fusarium oxysporum* f. sp. *pisi* (Landa et al. 2002) and black root rot of tobacco (Stutz et al. 1986). Previous work conducted in our laboratory showed the effectiveness of seed treatment with CHA0 to induce resistance against leaf rust caused by *Puccinia triticina* (unpublished). Also, in other studies, *P. fluorescens* species, including CHA0 and PCL, were reported to be efficient suppressive agents of fungal pathogens by inducing systemic resistance (Defago et al. 1990; Howell and Stipanovic 1980; Voisard et al. 1989; Tziros et al. 2007; Bardas et al. 2009). The control of *Z. tritici* by beneficial *P. fluorescens* was attributed to a direct inhibition *in situ* of the fungus by hydrogen cyanide and antimicrobial compounds (Flaishman et al. 1996; Levy et al. 1992). Also other biocontrol organisms such as *Bacillus subtilis* were effective in protecting wheat against STB disease through a direct antifungal activity of their cyclic lipopeptides (Mejri et al. 2018). In this study, inoculation of the STB-susceptible wheat cv. Spluga was performed using *Z. tritici* isolate 3D7, which was collected in a Swiss wheat field in 1999 and was found to be highly aggressive on several wheat cultivars (Zhan et al. 2002; Zhan et al. 2005). This high virulence might be the reason for the observed lack of resistance induction by the tested bacteria.

The present study shows that BABA applied as a soil-drench was effective in protecting wheat seedlings from *Z. tritici* infection, whereas, in plants soil-drenched with BTH, fungal growth was only delayed during the early germination phase. In this case, foliar application may be more effective, since BTH displayed a direct antifungal activity already at very low concentrations. Unexpectedly, wheat seed treatment with CHA0 or PCL did not enhance resistance to STB disease in wheat. Recently, Imperiali et al. (2017) demonstrated the possibility to combine CHA0 and PCL without affecting their capacity to colonize wheat roots. Hence, a combination of these two strains could result in a synergistic effect that may help to control STB disease, as was reported in other case of biological control in wheat (El-Sharkawy et al. 2018; Pierson and Weller 1994).

The results point to the possibility of developing effective protective measures against *Z. tritici* infection of wheat based on chemical inducer application. A histochemical analysis of the plant reactions during the infection process should provide more insights into the exact defense mechanism implicated in the observed resistance and allow for an adapted choice of inducer.

## Acknowledgements

We thank Daniel Croll, Leen Abraham and Nikhil Kumar Singh from the Evolutionary Genetics Laboratory, University of Neuchâtel, for technical support and advice to handle the STB pathogen. FB gratefully acknowledges the financial support by the Swiss Federal Commission for Scholarships for Foreign Students and BMM the financial support of the Swiss National Science Foundation, Grant No. 310030_160162).

## Supplementary material

**Table S1.** BABA levels in leaf tissues of wheat plants 2 and 6 days post tratement with BABA as soil-drench. The extraction and quatification of BABA was conducted as previously described by Thevenet et al. (2017).

| Treatment | BABA concentration (μM) |  |
| --- | --- | --- |
|  | 2 days | 6 days |
| H<sub>2</sub>O | 0.02 ± 0.009 | 0.02 ± 0.001 |
| BABA | 15.53 ± 4.95 | 4.58 ± 0.76 |

The values represent the mean ± SE of 3 biological replicates

